# Beyond acute toxicity: evolutionary response by rapid polygenic adaptation to a complex environmental stressor in *Chironomus riparius*

**DOI:** 10.1101/2025.05.23.655730

**Authors:** Lorenzo Rigano, Markus Schmitz, Henner Hollert, Markus Pfenninger

## Abstract

Anthropogenic stressors, such as pollution, habitat degradation, and climate change, are altering selective pressures on natural populations, but the evolutionary consequences of chronic exposure to complex mixtures of contaminants remain poorly understood. Addressing this knowledge gap is critical to the emerging field of evolutionary ecotoxicology, which aims to understand how long-term exposure to environmental contaminants shapes adaptive evolution and genome-wide variation. In this study, we employed urban runoff sediment as complex and environmentally realistic model stressor to investigate how multigenerational exposure affects fitness and potentially drives genomic adaptation in the freshwater midge *Chironomus riparius*. We combined an evolutionary life-cycle test with the Evolve and Resequence (E&R) approach, exposing replicate populations over seven generations to three treatments: an uncontaminated control and two concentrations of urban runoff sediment (0.5% and 10%). Key fitness traits, including mortality, mean emergence time (EmT50), fertility, and population growth rate (PGR), were measured, while allele frequency changes (AFC) were tracked to identify genomic signatures of selection. The results revealed distinct and non-linear fitness responses across treatments, including transgenerational effects, recovery of performance, and evidence of life-history trade-offs. Candidate haplotypes were enriched for genes involved in membrane transport, metabolism, and gene regulation, suggesting selection on general stress-response pathways consistent with polygenic adaptation. Signals of selection were also detected in control populations, underscoring the evolutionary influence of laboratory conditions. Overall, our findings demonstrate how evolutionary ecotoxicology can reveal both the potential and the constraints of rapid adaptation to realistic environmental stressors and highlight the importance of integrating evolutionary perspectives into ecological risk assessment.

## 1. Introduction

The increasing influence of anthropogenic stressors on the environment has reshaped selective pressures on natural populations, with pollutants, habitat destruction, and climate change recognized as key drivers of evolutionary change (Briski et al., 2025). These environmental pressures directly contribute to biodiversity loss and generate multifaceted selective forces that alter environmental conditions across multiple scales. As a result, exposed populations may undergo changes in key fitness traits, including growth, reproduction, and survival (Doria and Pfenninger, 2021; Rigano et al., 2024), ultimately restructuring population dynamics and ecosystem stability (McCann, 2000; Pennekamp et al., 2018).

Despite increasing recognition of chemical pollution as a major threat to biodiversity, research on its ecological and evolutionary consequences often remains compartmentalized. Ecotoxicological studies tend to focus on organismal-level effects or specific molecular mechanisms, while ecological research more commonly addresses large-scale patterns of biodiversity change, often without explicitly considering chemical stressors. This disciplinary separation has limited our ability to assess and predict pollution impacts at higher levels of biological organization. Bridging this gap requires integrative approaches that connect chemical exposure to molecular and phenotypic responses, and link these to ecological consequences at the population, community, and ecosystem levels across time and generations (Sylvester et al., 2023). In this context, the emerging field of evolutionary ecotoxicology combines ecotoxicology and evolutionary biology to understand how long-term exposure to pollutants influences evolutionary processes and shapes how natural populations adapt over time (Brady et al., 2017; Straub et al., 2020).

Adaptive responses can be observed as partial restoration of fitness, while their genetic basis can be inferred through genomic signatures of positive selection (Biswas and Akey, 2006). Positive selection occurs when beneficial alleles increase in frequency due to fitness advantages they confer, leaving detectable genomic patterns, such as changes in allele frequencies at and around selected loci (Manel et al., 2016; Nielsen, 2005). However, stressors in the environment, such as pollutants, rarely act in isolation. Instead, they interact in complex, non-linear ways, often generating unpredictable ecological and evolutionary responses (Rillig et al., 2019). Therefore, the interpretation of allele frequency change (AFC) can be challenging because simultaneous action of multiple selective pressures and non-adaptive processes, such as genetic drift, hitchhiking, and background selection, can hinder the association between a putative selective agent and the observed genomic response (Volis et al., 2005). In this context, integrating phenotypic and genomic responses becomes essential, as genomic data reveal the evolutionary potential and targets of selection, while life-history traits capture the realized fitness consequences.

Studies on single contaminants such as cadmium or microplastics have shown that evolutionary responses can vary widely, from partial fitness recovery to long-term population decline, depending on the type of stressor, exposure conditions, and genetic background of the affected population (Doria et al., 2025, 2021; Doria and Pfenninger, 2021; Khosrovyan et al., 2022). However, capturing ecologically realistic exposure scenarios means considering the simultaneous presence of multiple chemical stressors. In this study, we used sediments from urban runoff as such a scenario, reflecting a complex mixture of contaminants whose composition has been described in detail elsewhere (Rigano et al., 2025).

Urban runoff sediments accumulate in sedimentation basins that collect runoff from urban areas, particularly near high-traffic roads (Proteau et al., 2022). These sediments form complex matrices containing a wide range of contaminants, including tyre and road wear particles (TRWPs), heavy metals, microplastics and polycyclic aromatic hydrocarbons (PAHs) (Baptista Neto et al., 2016; Proteau et al., 2022; Rigano et al., 2025). Previous studies have shown that exposure to urban runoff sediments can increase mortality by up to 30%, reduce fertility, and decrease population growth rate (PGR) in *Chironomus riparius*, even after a single generation (Rigano et al., 2024). Chronic exposure over multiple generations has also been shown to increase germline mutation rates by approximately 50%, independent of concentration, with mutational spectra resembling those induced by Benzo[*a*]pyrene (BaP), a polycyclic aromatic hydrocarbon (PAH) (Rigano et al., 2025).

*Chironomus riparius*, a non-biting freshwater midge, has emerged as a key model in evolutionary ecotoxicology due to its short generation time, comprehensive genomic resources, and ecological relevance as a sentinel species in freshwater ecosystems (Foucault et al., 2019). Its sensitivity to environmental pollutants, combined with its ability to adapt to chemical stressors, makes it an ideal system for studying evolutionary responses to complex contaminant mixtures. Moreover, unlike many bioassays that use only sediment extracts or water-based samples, the *Chironomus* assay can be carried out with whole sediments, making it possible to assess toxicity in conditions that more closely reflect natural environments (OECD, 2023). As these previous studies have shown, *C. riparius*, as a model organism for freshwater biodiversity, may experience strong selective pressure from such complex environmental stressors, yet the long-term evolutionary consequences of such anthropogenic exposure remain largely unexplored.

To address this gap, experimental evolution combined with whole-genome sequencing offers a robust framework to investigate how populations adapt to environmental stressors across generations. In particular, the Evolve and Resequence (E&R) approach combines experimental evolution with high-throughput sequencing of pooled individuals (Pool-Seq) and has proven to be a reliable method for identifying genomic signatures of adaptation. In E&R studies, replicate populations are exposed to controlled selection pressures over multiple generations, and allele frequency changes are monitored over time to identify candidate loci under selection (Long et al., 2015; Pfenninger and Foucault, 2020; Schlötterer et al., 2015).

In this study, we used a multi-generational E&R approach with *C. riparius* exposed to urban runoff sediment to investigate: 1) How chronic exposure to this complex mixture impacts key fitness traits over multiple generations; 2) Whether genomic signatures of selection emerge in response to these conditions, and what is the nature of the adaptive changes at the genomic level; and 3) How phenotypic and genomic responses co-evolve over time under this ecologically relevant stressor.

## 2. Materials and Methods

### 2.1 Sediment collection, preparation and composition

In February 2023, sediment samples were collected from an urban runoff sedimentation basin located in Aachen, Germany (50°48’03.6”N 6°06’29.7”E). This basin receives runoff via a separate sewer system from a large area in the district of Aachen Soers, including a sports park and a heavily trafficked road, Bundesstraße B57 Krefelder Straße. The total drained area covers approximately 68 hectares. The B57 serves around 26,000 vehicles per day, based on a 2021 traffic count by the Federal Highway Research Institute (BASt). Sampling was conducted during routine cleaning of the basin. Sediment was collected from multiple locations within the basin using a stainless-steel bucket, pooled to form a composite sample, and manually homogenized. The homogenized sample was stored in a cold room at 4 °C until further processing, which was completed in the following weeks. For subsequent analysis, aliquots were freeze-dried and sieved using a stainless steel mesh (1 mm) to remove large particles (Shuliakevich et al., 2022).

The sediment used in this study corresponds to the same matrix previously characterized in detail by Rigano et al., (2025). The urban runoff sediment contained a broad spectrum of urban and traffic-related contaminants, including tyre rubber particles (up to 5.2 g/kg), heavy metals (e.g., Fe: 19.5 g/kg, Zn: 1.74 g/kg, Cu: 144 mg/kg), road salts (46.12 g/kg), and mineral oils (983 µg/kg). The total concentration of PAHs was 4.0 mg/kg, with high molecular weight compounds such as Benzo[*b*]fluoranthene and Benzo[*g,h,i*]perylene accounting for approximately 80% of the total. LC-HRMS/MS screening further detected 124 synthetic organic compounds, including tyre additives (e.g., phosphate esters, phthalates, benzothiazoles, bisphenols), as well as personal care products, pesticides, biocides, and PFAS. A comprehensive chemical profile is available in Rigano et al., (2025).

### 2.2 Source population and establishment of laboratory population

The *C. riparius* culture used in this study originated from a local population collected from a small river in Hasselbach, Hessen, Germany (50.167562N, 9.083542E). The culture was maintained as an in-house laboratory standard and regularly supplemented with individuals from the field to preserve genetic diversity. Stock culture conditions followed a modified protocol based on OECD guideline No. 219 (Foucault et al., 2019).

### 2.3 Evolutionary life-cycle test

An uncontaminated control and two treatment groups exposed to urban runoff sediment (0.5% and 10% by total weight) were tested. The control group consisted of 500 g of quartz playground sand. The 0.5% group contained 497.5 g of quartz sand mixed with 2.5 g of urban runoff sediment, while the 10% group consisted of 450 g of sand and 50 g of sediment.

To initiate the life-cycle test in each generation, 10 freshly laid egg ropes were collected at the beginning of each generation from the same E&R replicates (see below) and transferred to 6-well plates containing 3 mL of medium per well. Nine glass bowls (Ø 20 × 10 cm) were prepared one day prior to the start of the experiment: three were assigned to the control group, three to the 0.5% group, and three to the 10% group.

Each bowl was filled with 1.25 L of medium and aerated continuously, except for the first 6 hours to allow larvae to settle in the sediment. Sixty first-instar larvae were introduced into each bowl at the beginning of each generation. The test was conducted over a period of 28 days, approximately corresponding to the duration of one generation, in accordance with OECD Guideline No. 233. (OECD, 2010). The organisms were fed daily with finely ground Tetramin® Flakes, with feeding amounts adjusted according to developmental stage (Foucault et al., 2019).

Adult emergence was recorded daily to calculate mortality by subtracting the number of emerged adults from the initial number of exposed larvae. The sex of emerged adults was recorded daily to calculate the mean emergence time (EmT50), defined as the time when 50% of the females had emerged. The number of fertile egg ropes was recorded, and egg counts were determined (Foucault et al., 2019). Finally, all measured parameters were summarized into PGR (Nemec et al., 2013). Data for the sixth generation of the control group could not be collected due to technical limitations.

Raw data were entered, and fitness parameters were calculated using Microsoft Excel® (MS Excel®). Statistical and graphical analyses were performed in R, applying linear, quadratic (second-order polynomial), and constant models to assess patterns in fitness parameters across contaminant exposure levels over multiple generations.

### 2.4 Establishment of the test replicates for E&R

Twenty-five egg ropes from the laboratory population were collected and hatched, and the ∼ 15,000 larvae pooled to establish nine experimental replicates. From these pools, at least 1,000 first-instar larvae were transferred to glass bowls (Ø20 × 14.5 cm) to initiate each replicate. Three replicates were assigned to each treatment group (0, 0.5%, 10%).

All replicates were maintained at 23°C, 60% humidity, and a 16:8 light:dark photoperiod, following OECD guideline No. 233 (OECD, 2010). The quartz playground sand was pH-neutral and washed before use. Urban runoff sediment and sand were mixed thoroughly using a laboratory spatula to homogenize the sediment column. The bowls were filled with 1.5 L of medium consisting of deionized water adjusted to a conductivity of 550–650 µS/cm with aquarium sea salt (TropicMarin®) and a pH of ∼8. Water loss due to evaporation was compensated by adding deionized water. The organisms were kept in swarming cages (40 × 40 × 60 cm) for the entire E&R experiment and fed weekly with finely ground Tetramin® fish flakes. Replicates were maintained under these consistent conditions for seven generations (Foucault et al., 2019).

### 2.5 Pool-seq-based detection of selection and haplotype analysis

Every 28 days, corresponding to the generation length at the experimental temperature, 120 adults were collected and pooled from each replicate over seven generations. In the F0 group, pooling was performed only at the end of the first generation, resulting in a total of 64 pools, including the F0 group and three replicates for each treatment: control (0), 0.5%, and 10%.

DNA was extracted from pooled samples of 120 individuals from the F0 and each replicate using the Qiagen blood and tissue extraction kit. Sequencing was performed on the Illumina NovaSeq platform, with an effective mean coverage ranging between 30× and 85×. Adapters and low-quality reads were trimmed using Trimmomatic (Bolger et al., 2014). Clean reads were then mapped to the latest *C. riparius* reference genome (v.4, unpublished) using the BWA mem algorithm (unpublished data) (Li and Durbin, 2009). Low-quality reads were filtered, and single nucleotide polymorphisms (SNPs) were initially called using samtools (Li et al., 2009).

The softwares Popoolation and Popoolation2 (Kofler et al., 2011a, 2011b) were used to call SNPs, remove indels, and estimate genetic diversity as Watterson’s theta in the first and last generations of each replicate. A synchronized file was generated using PoPoolation2 and processed with a custom Python script to estimate allele frequencies. The sync file was filtered using the following parameters: minimum SNP coverage (≥ 15), minimum read count per population (≥ 15), minor allele frequency (≥ 0.10), and variance among allele frequencies (≥ 0.10).

Potentially selected SNPs were identified by detecting AFC larger than expected under genetic drift. AFC across generations were assessed using two complementary statistical approaches: linear regression, to detect SNPs exhibiting consistent frequency shifts over time, and Fisher’s exact test, to identify SNPs showing significant differentiation between the initial and final generations. The top 1% of outliers from both tests were cross-referenced to identify selection signals.

To account for the expected effect of genetic drift, replicate-specific effective population sizes (N_e_) were first estimated from the observed AFC between F0 and the final generation. Expected allele frequency trajectories under drift were then simulated using the Wright-Fisher model with the estimated N_e_ values over seven generations. Significance of the observed allele frequency changes was tested using a chi-square test, followed by the Benjamini-Hochberg (BH), comparing observed and expected values under drift. A 99.9th percentile threshold was used to define highly significant SNPs according to the chi-square test and to compare real observations with expected drift. Only SNPs with -log10(p-value) exceeding the 99.9th percentile of the simulated distribution were retained for further analysis. All calculations and simulations were performed with the R package *PoolSeq* (Taus et al., 2017).

The number of candidate SNPs was likely inflated by the hitchhiking of neutral variants linked to selected SNPs, driven by the low number of recombination events expected during the experiment, which increased the frequency of large haplotype blocks (Franssen et al., 2017). To identify haplotypes, potential marker SNPs were defined as those falling within the top 20th percentile of AFC values. Pearson correlation coefficients were calculated between each marker SNP and neighbouring SNPs to define haplotypes boundaries. Haplotype blocks were extended from each marker SNP until the correlation coefficient with the focal marker fell below that with the next neighbouring SNP. If the correlation with the next marker exceeded that with the focal marker, the block was discontinued, and a new haplotype was initiated. Haplotypes were considered to overlap within and between treatment groups if they extended at least 25% in both directions from the marker SNP. After haplotype assignment, a Monte Carlo intersection analysis was performed to assess whether their presence within and between treatments aligned with random expectations.

The selection coefficient (s) was estimated from the change in allele frequency over the course of the experiment for each selected SNP (Crow, 2000). For each haplotype, the SNP with the highest selection coefficient was identified as the most likely selection target. Although a single primary selection target may exist within each haplotype, the entire haplotype with all its phenotype-relevant variants is likely under selection.

The gene nearest to the SNP with the highest selection coefficient within each haplotype was tentatively considered a potential selection target across all replicates. The resulting list of candidate genes was analysed for overrepresentation of gene ontology (GO) terms related to “biological function” using Fisher’s exact test implemented in the R package TopGO (Alexa, Adrian, and Jörg Rahnenführer, 2009).

## 3 Results

### 3.1 Evolutionary life-cycle test

Mortality, mean emergence time (EmT50), fertility, and PGR exhibited distinct patterns across the control, 0.5%, and 10% groups over seven generations. In the control, mortality increased steadily over generations, following a linear model (R^2^ = 0.277, p = 0.0248), whereas in the 0.5% group, mortality increased in a polynomial fashion (R^2^ = 0.251, p = 0.0464). In contrast, mortality in the 10% group followed a quadratic pattern (R^2^ = 0.444, p = 0.2177), with an initial increase followed by partial stabilization in later generations (Figure 1a).

**Figure 1.**
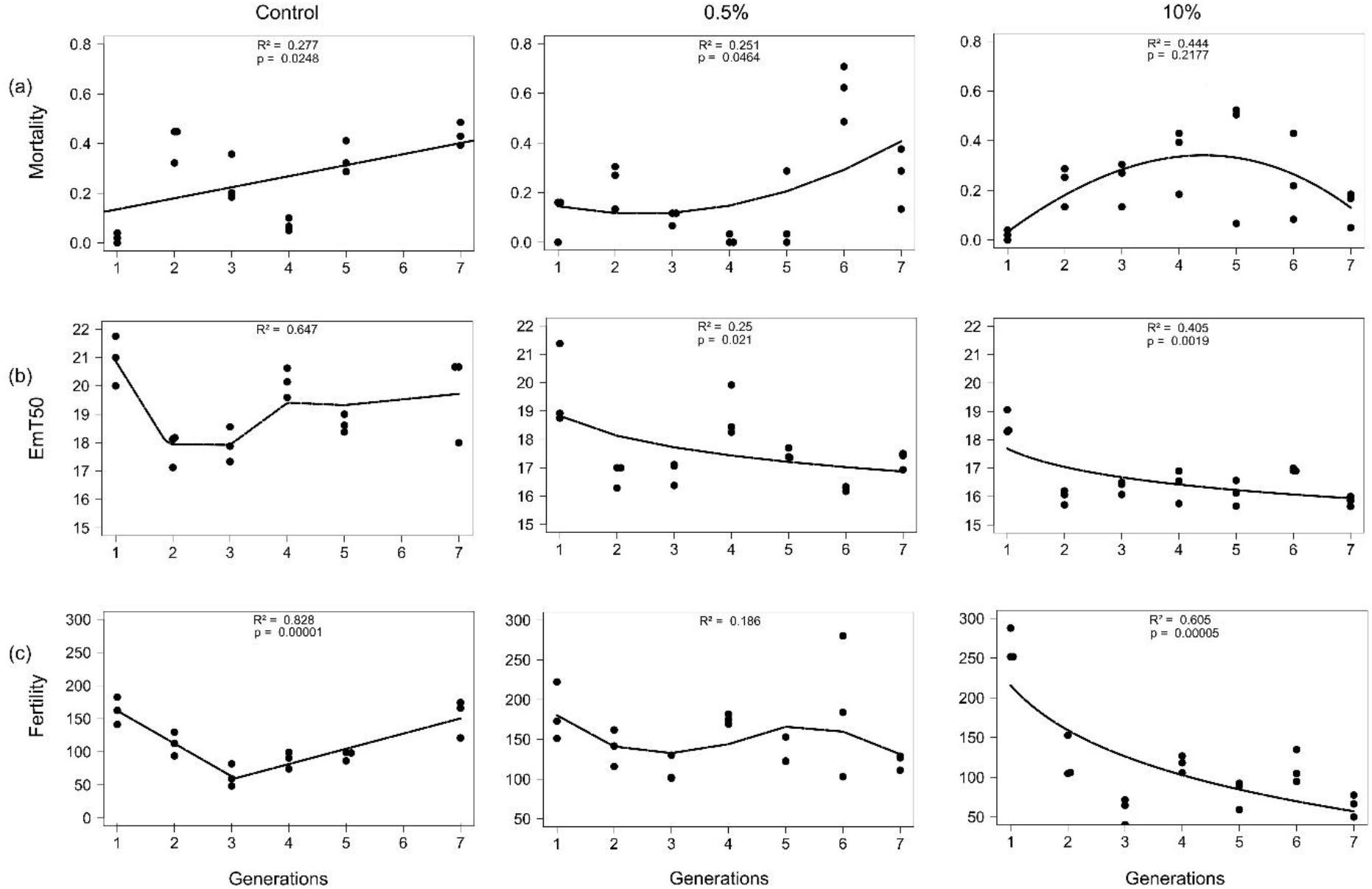
*C. riparius* life-cycle endpoints measured over seven generations: (a) Mortality, (b) EmT50, and (c) Fertility. Each data point represents the value from an independent replicate population, based on multiple individuals per generation and treatment. Trend lines represent the best-fitting models for each treatment and endpoint, selected based on visual inspection and model fit (R^2^), and include linear, polynomial, quadratic, logarithmic, segmented linear, and loess models. Loess models were used for descriptive smoothing and do not yield standard p-values.

EmT50 exhibited contrasting patterns across treatments (Figure 1b). In the control, EmT50 showed a slight, non-significant decrease over generations (R^2^ = 0.647), following a loess model. In the 0.5% group, EmT50 declined significantly (R^2^ = 0.25, p = 0.021), following a logarithmic model, while the most pronounced decline was observed in the 10% group (R^2^ = 0.405, p = 0.0019), following a power model.

Fertility decreased over generations in the control, following a segmented linear pattern, but showed recovery after the third generation (R^2^ = 0.828, p = 0.00001). In the 0.5% group, fertility declined with no significant trend (R^2^ = 0.186). In contrast, fertility in the 10% group decreased significantly over generations (R^2^ = 0.605, p = 0.00005) (Figure 1d).

PGR in the control exhibited a significant downward trend, following a segmented linear model (R^2^ = 0.655, p = 0.0006), with a decline up to the third generation, followed by stabilization in subsequent generations. In the 0.5% group, no significant change in PGR was observed (R^2^ = 0.201). In the 10% group, PGR showed a similar trend to the control, declining significantly (R^2^ = 0.613, p = 0.0485), but the decrease was less pronounced after the third generation, followed by partial recovery (Figure 2a).

**Figure 2.**
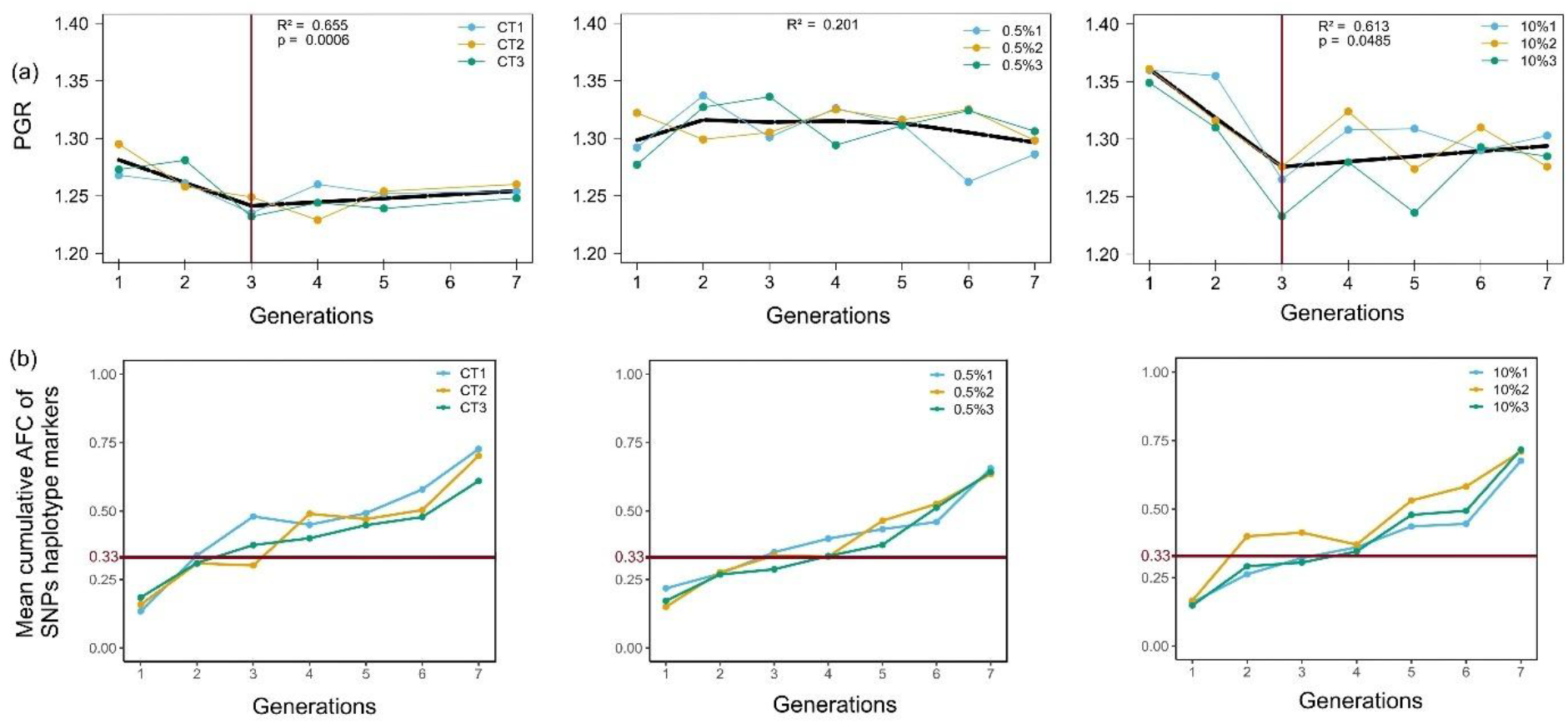
(a) PGR measured across seven generations for each replicate. Red vertical lines indicate the generation at which the inflection point in the PGR trajectory was observed. (b) Mean cumulative AFC of haplotype-defining SNPs per replicate. The horizontal red line marks the upper 95% confidence limit expected under genetic drift.

### 3.2 Pool-seq-based detection of selection and haplotype analysis

The raw sequencing data and pool mapping files are available at the European Nucleotide Archive (ENA), under accession number PRJEB89250.

The replicate-specific effective population size (Ne) during the experiment varied between 29 and 115 (mean ± SD: 63.75 ± 27.18) (Table 1). The F0 population had a mean genome-wide population mutation parameter theta of 0.012. Based on the species’ point mutation rate of 2.1 × 10^−9^ per site per generation (Oppold and Pfenninger, 2017), this corresponds to a long-term N_e_ of approximately 1.43 million. In the evolved replicates, theta ranged between 0.005 and 0.012. Selection and drift jointly reduced genetic variation by up to 53.1%, although in some replicates the reduction was negligible compared to the F0 population (Table 1).

**Table 1:** Demographic and genetic parameters of each evolved replicate population. Reported are the estimated effective population size (Ne), the percentage loss of nucleotide diversity relative to the ancestral (F0) population (Genetic reduction %), the average AFC of selected SNPs, and the number of haplotypes detected.

| Replicate | $N_e$ | Genetic reduction (%) | AFC of sel. SNPs | N. haplotypes |
| --- | --- | --- | --- | --- |
| Control1 | 28.75 | 6.81 | 0.72 | 39 |
| Control2 | 40.42 | 8.60 | 0.70 | 44 |
| Control3 | 71 | 17.02 | 0.61 | 44 |
| 05%1 | 89.72 | 10.27 | 0.65 | 25 |
| 05%2 | 77.79 | 52.61 | 0.63 | 56 |
| 05%3 | 115.24 | 31.55 | 0.64 | 17 |
| 10%1 | 53.66 | 8.35 | 0.67 | 34 |
| 10%2 | 47.78 | 8.74 | 0.71 | 27 |
| 10%3 | 49.35 | 2.61 | 0.71 | 32 |

The average AFC per replicate, calculated between the F0 and F7 generations for the selected SNPs, ranged between 0.61 and 0.72 (Table 1). A total of 1,171 SNPs in the control, 538 in the 0.5% urban runoff sediment treatment, and 674 in the 10% urban runoff sediment treatment exceeded the expected threshold and were retained for haplotype analysis. These SNPs allowed the identification of 318 haplotypes: 127 in the control replicates, 98 in the 0.5% urban runoff sediment replicates, and 93 in the 10% urban runoff sediment replicates (Table 1). An ANOVA followed by a Tukey post-hoc test showed significant differences in haplotype numbers only for two replicates of the 0.5% urban runoff sediment treatment (p = 0.025). A Kruskal-Wallis test followed by a Dunn post-hoc test with Bonferroni correction also showed significant differences in haplotype length between the same two replicates (p = 0.014). Haplotypes were considered overlapping if their lengths, extending in both directions from the marker SNP, overlapped by at least 25%. Based on this criterion, 9 haplotypes were assigned to the control, 1 to the 0.5% treatment, and 2 to the 10% treatment. Additionally, 12 haplotypes were shared between the control and 0.5%, 17 between the control and 10%, 15 between 0.5% and 10%, and 8 were shared among all three treatments (Figure 3b).

**Figure 3.**
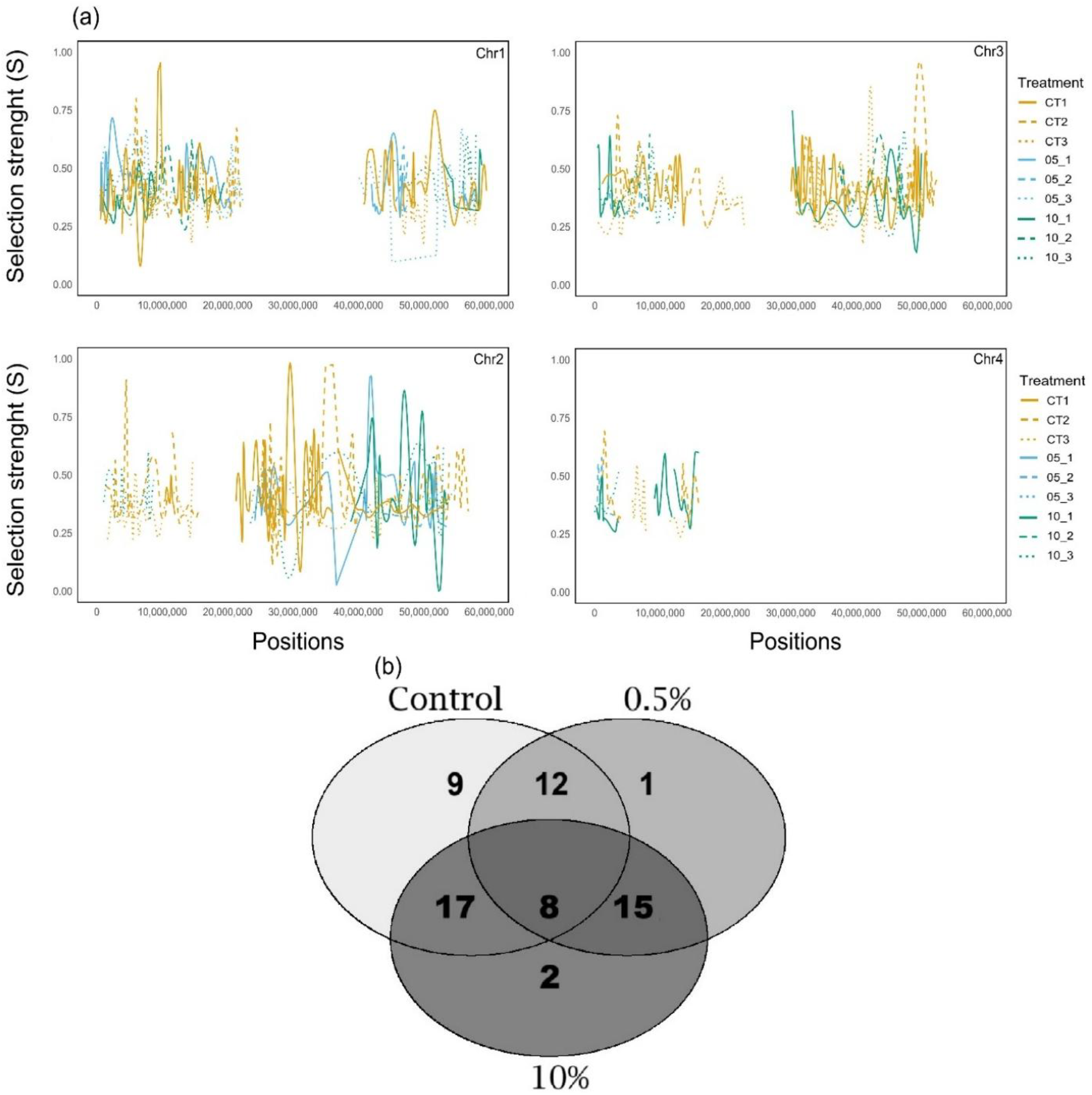
(a) Distribution of haplotypes across chromosomes in the different treatments. (b) Venn diagram showing the number of exclusive and shared haplotypes among treatments.

The Monte Carlo intersection analysis revealed that the number of observed haplotypes largely aligned with the expected random distribution, with some deviations depending on the treatment (Table 2). In the control group, the number of observed haplotypes (9) was very close to the expected value (8.63) and showed no significant deviation (p = 0.89).

The degree of haplotype sharing between treatments showed more pronounced differences. The number of haplotypes shared between the control and 0.5% groups (12) was higher than expected (7.22) but did not reach statistical significance (p = 0.058). In contrast, haplotype sharing between the control and 10% groups (17) and between the 0.5% and 10% groups (15) was significantly higher than expected (p = 0.003 and p = 0.0003, respectively). The overlap between all three treatments (8) was not significantly different from the expected value (6.54) (p = 0.548). In contrast, the number of haplotypes in the 0.5% and 10% treatments (1 and 2, respectively) was lower than expected (7.01 and 9.83), and this reduction was statistically significant in both groups (p = 0.016 and p = 0.006).

Estimated selection coefficients for significant SNPs ranged between 0.35 and 0.45. The GO term enrichment analysis identified several biological processes significantly overrepresented among the genes closest to SNPs within shared haplotypes across treatments. In haplotypes shared between the control and 0.5% groups, the most enriched terms were linked to dipeptide import across the plasma membrane (GO:0140206; p = 0.00000018), unsaturated fatty acid biosynthetic process (GO:0006636; p = 0.00003454), and synapse organization (GO:0050808; p = 0.00018).

Haplotypes shared between the control and 10% groups showed enrichment for processes such as anion transport (GO:0006820; p = 0.00000016), sulfation (GO:0051923; p = 0.00000898), and proteolysis (GO:0006508; p = 0.00005742).

For haplotypes shared between the 0.5% and 10% groups, the most enriched terms included NAD-cap decapping (GO:0110155; p = 0.00000004), O-glycan processing (GO:0016267; p = 0.000000065), and proteolysis (GO:0006508; p = 0.000007469), suggesting selective pressure on post-transcriptional and protein turnover pathways. Strong enrichment was also observed for nucleic acid phosphodiester bond hydrolysis (GO:0090305; p = 0.000016749) and regulation of transcription by RNA polymerase II (GO:0006357; p = 0.000086369), indicating targeted selection on gene expression and nucleic acid metabolism.

## 4 Discussion

Environmental contaminants are well known to impair fitness and alter the genomic architecture of exposed organisms. However, it remains poorly understood whether these ubiquitous stressorscan act as selective agents and drive adaptive evolution, especially when occurring in complex mixtures. To address this, we employed a multigenerational evolution approach that integrates life-history data with the E&R framework. This combined design allowed us to simultaneously assess phenotypic responses and genome-wide AFC in *C. riparius* populations exposed over seven generations to different concentrations of urban runoff sediments.

Our study provides one of the first integrative assessments of both adaptive and mutational responses to a realistic contaminant mixture over multiple generations, offering novel insight into the evolutionary dynamics triggered by chronic pollutant exposure.

### 4.1 Phenotypic and genomic responses to multigenerational exposure

To assess the effects of chronic exposure on fitness, we monitored mortality, EmT50, fertility and PGR. Phenotypic responses revealed nonlinear trajectories and potential trade-offs, suggesting that adaptive changes do not scale linearly with contaminant concentration. These patterns were accompanied by detectable changes in allele frequency, discussed below, indicating that the observed phenotypic dynamics were underpinned by genomic responses. We also detected evidence of selection in the control group, indicating that laboratory conditions imposed selective pressures, highlighting the importance of distinguishing these effects from those caused by contaminant exposure.

In the control group, mortality progressively increased over successive generations, likely due to nutritional limitations or rearing conditions differing from those of the ancestral stock culture. The temporal consistency of this pattern supports the hypothesis of a sustained physiological burden, rather than random demographic fluctuations. Conversely, the initially low mortality in the 0.5% group may reflect a nutritional benefit from organic content in the sediment, temporarily offsetting toxic effects (Rigano et al., 2024). However, this benefit did not prevent long-term physiological compromise, as mortality increased over time, highlighting that even low-level exposure can lead to significant fitness costs. In the 10% group, mortality exhibited a biphasic trajectory, with an initial peak, presumably caused by acute toxicity, followed by partial stabilization in subsequent generations. Therefore, in contrast to the lower concentration, the high contaminant load in the 10% group likely neutralized any potential nutritional advantage, with toxicity emerging as the dominant driver of phenotypic responses.

EmT50 was also influenced by treatment, reflecting stress-induced developmental shifts. In the control group, EmT50 initially decreased and later recovered. In contrast, urban runoff sediment-exposed individuals, particularly at 10%, exhibited accelerated development across generations. This may be linked to increased nutrient availability in the urban runoff sediment, consistent with findings by Rigano et al. (2024), who reported accelerated development and increased larval size after a single generation of exposure to a similar matrix. However, faster development may also represent a stress-induced life-history adjustment. Kolbenschlag et al. (2024) showed that exposure to *Bacillus thuringiensis israelensis* (Bti), a biological larvicide, induced earlier emergence in *C. riparius*, together with morphological alterations and reduced adult fitness. These findings suggest that accelerated development may arise from distinct environmental pressures and involve physiological trade-offs, such as reduced energy reserves or compromised reproductive capacity (Zera and Harshman, 2001).

In the control group, fertility declined during generations F2 and F3 and then recovered from F4 onwards, reflecting an initial response to stress imposed by laboratory conditions, followed by recovery in subsequent generations. The 0.5% group maintained a relatively stable fertility across generations, suggesting that this exposure level did not significantly impair reproductive function, at least within the timeframe of the experiment. In contrast, fertility steadily declined in the 10% group, consistent with a likely cumulative reproductive cost associated with sustained contaminant pressure.

The observed fitness dynamics became particularly evident when considering PGR, a composite measure that integrates all the fitness parameters. In both the control and 10% groups, PGR declined during generations F2–F3, followed by a recovery in subsequent generations. In contrast, the 0.5% group showed no consistent deviation in PGR, suggesting that this lower exposure level remained below the threshold required to elicit long-term effects.

The similarity in PGR dynamics between the control and 10% groups suggests a shared underlying mechanism. This segmented pattern contrasts with theoretical expectations, which predict that fitness responses to environmental stress occur rapidly, usually within one generation, as selection quickly favours traits improving survival and reproduction (Fisk et al., 2007; Goitom et al., 2018; Sibly and Calow, 1989). We hypothesise that stress experienced by females in the early F1 generation compromised resource allocation into reproduction, thereby diminishing egg rope quality. This might have delayed visible fitness restoration in subsequent generations, despite the immediate onset of molecular selection (see below). These types of transgenerational effects, although often transient, have been documented in both invertebrate and vertebrate models. For instance, Salesa et al. (2022) reported that *Daphnia magna* exposed to prochloraz exhibited no reproductive effects in the F0 generation but showed significant phenotypic changes in F1. Similarly, exposure of pregnant rats to endocrine disruptors impaired the fertility of male offspring up to the F4 generation (Anway et al., 2005).

Although allele frequency changes were detectable from the earliest generations, phenotypic recovery did not occur immediately, suggesting that beneficial alleles required time to rise sufficiently in frequency to produce a measurable effect on population fitness. Suggestively, from generation F3 onward, allele frequency changes at haplotype-defining SNPs exceeded expectations under genetic drift across all treatments. Thus, the initial decline and subsequent recovery of PGR from generation F4 onward in the control and 10% groups may reflect the interaction between transgenerational physiological constraints and evolutionary adaptation.

Genetic drift can strongly affect small experimental populations (Lynch et al., 2016). Additionally, limited recombination and a small number of generations constrain the breakdown of linkage disequilibrium (LD), causing selected alleles to remain physically linked to nearby neutral variants and hindering the identification of loci directly under selection (Kofler and Schlötterer, 2014). Although the E&R approach is limited in short-term experiments, due to low recombination and small population sizes that can reduce resolution and hinder causal loci (Franssen et al., 2017; Otte and Schlötterer, 2021), the consistent deviations from drift expectations across independent replicates, along with the functional enrichment of selected regions, support the conclusion that selection drove the observed genomic changes. Moreover, the temporal correspondence between molecular shifts and phenotypic recovery aligns with models of polygenic adaptation, in which selection acts on multiple small-effect loci distributed throughout the genome (Barghi et al., 2020; Berg and Coop, 2014).

These selective responses, despite differences in phenotypic outcomes across treatments, revealed shared genomic patterns. Functional annotation of candidate regions indicated that selected haplotypes were significantly enriched for biological processes involved in membrane transport, lipid metabolism, proteolysis, and gene expression regulation. This suggests that adaptation predominantly targets general stress-response pathways aimed at maintaining cellular homeostasis under chronic exposure, rather than pathways specific to contaminant concentration. These observations align with models of polygenic adaptation, in which selection acts on standing genetic variation across broad regulatory networks (Barghi et al., 2020, 2019).

### 4.2 Evolutionary implications and ecological relevance

Our study revealed coordinated phenotypic and genomic responses in both the control and 10% treatments. While the changes observed in the control group likely reflect adaptation to laboratory conditions, the common response in the sediment treatments suggested that chronic exposure to urban runoff sediment imposed selective pressures that contributed to evolutionary change. Similar patterns have been reported in previous multigenerational experiments involving single contaminants, such as heavy metals and microplastics, which are also major constituents of the complex mixture investigated in this study. For example, *C. riparius* exposed to increasing cadmium concentrations over eight generations exhibited genome-wide allele frequency changes indicative of strong selection, despite only modest phenotypic changes (Doria et al., 2022). Likewise, exposure to polyamide microplastics initially caused high larval mortality, followed by a rapid compensatory response and substantial allele frequency changes, consistent with polygenic adaptation (Khosrovyan et al., 2022). In another organism, *Daphnia magna* exposed to perfluorooctane sulfonate (PFOS) experienced reduced individual fitness during chronic exposure, while population growth recovered over successive generations, suggesting a potential for adaptive compensation despite sustained physiological stress (Tae-Yong Jeong et al., 2016).

Beyond selection, prior work using the same sediment matrix showed a significant increase in germline mutation rate in *C. riparius*, by approximately 50%, regardless of concentration (Rigano et al., 2025). This highlights a dual role of urban runoff sediment in shaping evolutionary trajectories, acting both as a selective pressure that promotes adaptive change and as a source of mutational input that increases genetic variation. This combination may accelerate evolutionary dynamics but also raises concerns about long-term genomic stability and fitness in natural populations.

By linking phenotypic trajectories with genome-wide allele frequency dynamics across generations, our study provides an integrated perspective on how aquatic organisms respond to sustained environmental stress. Although conducted under controlled laboratory conditions with relatively small populations, we detected clear adaptive responses at both the phenotypic and genomic levels. This suggests that natural populations, typically characterized by greater effective population size and thus genetic diversity and subjected to continuous exposure, may also evolve in response to similarly complex pollutant mixtures. However, the increased ecological complexity and co-occurrence of multiple stressors in natural environments are likely to influence both the direction and pace of such adaptation (Rullens et al., 2022).

In summary, our findings underscore the urgent need to integrate evolutionary perspectives into environmental risk assessment and regulatory frameworks. As pollution continues to reshape the genetic and ecological fabric of natural populations, ecotoxicology must move beyond short-term toxicity endpoints to consider the long-term evolutionary consequences of chronic exposure. Understanding how contaminants alter not only survival but also the heritable basis of adaptation is essential to safeguard biodiversity and ecosystem stability. In an era of accelerating environmental change, failing to do so risks underestimating the true cost of pollution, for nature and for society.

## Supporting information

Supplementary Infos

## Acknowledgments

The study received support from the LOEWE-Centre Translational Biodiversity Genomics (TBG) (LOEWE/1/10/519/03/03.001(0014)/52) funded by Hessen State Ministry of Higher Education, Research and the Arts (HMWK). The RoadTox project was funded by the Ministry of the Environment, Nature and Transport of the State of North Rhine-Westphalia and the RobustNature Excellence network by internal funding of Goethe-University, Frankfurt.

